# Multi view based imaging genetics analysis on Parkinson disease

**DOI:** 10.1101/2021.08.19.456943

**Authors:** Guglielmo Cerri, Manuel Tognon, Simone Avesani, Neil P. Oxtoby, Andre Altmann, Rosalba Giugno

## Abstract

Longitudinal studies integrating imaging and genetic data have recently become widespread among bioinformatics researchers. Combining such heterogeneous data allows a better understanding of complex diseases origins and causes. Through a multi-view based workflow proposal, we show the common steps and tools used in imaging genetics analysis, interpolating genotyping, neuroimaging and transcriptomic data. We describe the advantages of existing methods to analyze heterogeneous datasets, using Parkinson’s Disease (PD) as a case study. Parkinson’s disease is associated with both genetic and neuroimaging factors, however such imaging genetics associations are at an early investigation stage. Therefore it is desirable to have a free and open source workflow that integrates different analysis flows in order to recover potential genetic biomarkers in PD, as in other complex diseases.

## 1 Introduction

Imaging genetics is an emerging field that bridges genetic insights into the biology of complex diseases and behaviors with rich, quantitatively observed imaging phenotypes. The primary goals of imaging genetics are to identify and to characterize genes influencing neuroanatomical or neurophysiological variation using brain images [1, 2].

The establishment of the ENIGMA consortium [3, 4] in 2009 significantly improved quality and reliability of *imaging genetics* studies. By rallying researchers from different fields, such as genetics, neurology, psychology, statistics and machine learning, ENIGMA aims to push forward *imaging genetics* to characterize brain structure and functioning by incorporating neuroimaging and genomic data. In [5], Richiardi and colleagues exploited *imaging genetics* power to study brain activity during rest in healthy adolescents. By combining gene expression data and functional MRI data, which measure brain activity through changes in blood flow, the authors showed that the co-expression pattern of a set of 136 genes affects the resting-state functional connectivity of the brain.

Moreover, *imaging genetics* analyses have been widely applied to characterize complex traits, such as neurodegenerative diseases. Lorenzi and colleagues [6] proposed a Partial Least Square model (PLS) to characterize the joint variation of genotype and phenotype in Alzheimer’s disease (AD) patients. To train the PLS model, the authors combined genetic and neuroimaging data, which represented the phenotypic information, discovering several Single Nucleotide Polymorphism (SNPs) that were statistically associated with AD. By combining the PLS model with functional prioritization [7], Lorenzi and colleagues found several polymorphisms involved in the cortical thinning observed in AD patients, by testing the candidate SNPs against expression Quantitative Trait Loci data [8]. The authors demonstrated a link between tribbles pseudokinase 3 (*TRIB3*) and the stereotypical pattern of gray matter loss in AD. *Imaging genetics* longitudinal analysis, involving genotyping and neuroimaging data, were also performed through Genome Wide Association studies (GWAS). In [9], by performing a GWA study using T1-weighted MRI and amyloid PET images as quantitative phenotypic traits, Scelsi and coworkers identified a polymorphism mapping within *LCORL*, a gene linked with AD and Parkinson’s disease (PD). Beside the important biological discovery, the authors demonstrated that multimodal imaging phenotypes may assist in unveiling genetic drivers related to the onset and progression of complex traits.

Despite the success in characterizing complex neurodegenerative diseases, such as AD, *imaging genetics* analyses have been scarcely applied on PD. However, studies combining neuroimaging and genomic data have been shown effective in discovering potential genetic drivers of PD [10, 11]. Kim and coworkers investigated the factors underlying the Movement Disorder Society-sponsored unified Parkinson’s disease rating scale (MDS-UPDRS) scores in PD patients [10]. The authors showed that linear models trained on genetic and neuroimaging data improve predictions on MDS-UPDRS scores, suggesting that genomic and brain imaging data provide complementary information in PD. In [11], Kim and colleagues proposed a joint connectivity-based sparse canonical correlation analysis (JCB-SCCA) combining genetic and neuroimaging data through an association analysis between SNP and multimodal neuroimaging data. By using genotyping, dMRI and DaT-SPECT data, the authors reported several genetic variants and brain regions potentially associated with PD development.

In this work we review the key steps of *imaging genetics* analysis by using PD-related genetic variants, gene expressions and imaging data, collected from the PPMI data portal [12]. We propose a multi view-based workflow to integrate multi-source data and provide a holistic functional interpretation of the candidate genetic variants which could constitute potential biomarkers for PD onset and progression. The proposed method covers the key steps of an *imaging genetics* analysis, from data preprocessing to results interpretation, integrating genetic, brain imaging and transcriptomic data. The multi view-based workflow is composed of three main phases: *Individual view, Integrated view* and *Functional interpretation*. The *Individual view* focuses on finding potential genetic biomarkers by interpolating genotyping and neuroimaging data, considering phenotypic traits individually. *Integrated view* phase is aimed to search for genetic variants-phenotype associations, summarizing different traits in a single model.m The *Functional interpretation* step focuses on linking the previously obtained results with their potential functional consequences. The source code of the multi view based workflow is freely distributed to the communities as indicated in the Data availability section.

## 2 Materials and Methods

In this section we first describe the data used throughout the manuscript, then we review the current state-of-the-art methods for imaging genetics analyses through the proposal of a multi-view-based workflow.

### 2.1 Data

Data used in the manuscript were retrieved from PPMI consortium data portal [12] (data downloaded in March 2020 via the PPMI website, http://www.ppmi-info.org/). The PPMI consortium focuses the efforts in identifying new potential biomarkers for PD progression and onset. Moreover, PPMI aims to enhance the development of new therapies and treatments for PD through longitudinal studies considering different types of data.

Throughout this study, we use genetic (genotyping and transcriptomic) and neuroimaging (DaTscan and MRI) data. We selected DaTscan and MRI data since they have been both shown to be proficient potential biomarkers for PD onset and progression [13–16], and represent different imaging information levels.

Genotyping data provides a set of DNA sequence polymorphisms (SNPs and indels). We used two different genotyping datasets in our study: ImmunoChip [17] and NeuroX [18, 19]. The ImmunoChip targets genetic loci that are known to be associated with major autoimmune and neuroinflammatory diseases [17]. The ImmunoChip dataset provides 196,524 genetic variants: 718 indels and 195,806 SNPs, with 1920 SNPs replicating PD associated genetic loci. ImmunoChip genotypes were obtained from 523 subjects. The NeuroX genotyping array targets ~240,000 exonic genomic variants and ~24,000 custom variants, involved in neurological diseases [18, 19]. NeuroX genotyping data are available for 619 subjects.

PPMI transcriptomic data measure the amount of RNA produced by transcription of each gene, with the aim of providing a comprehensive view of the role played by gene expression and biological pathways in PD onset and progression. Transcriptomic data were retrieved by performing a whole transcriptome RNA-seq experiment on whole blood samples. Whole transcriptome sequencing was sequenced at Hudson Alpha’s Genomic Services Lab on an Illumina NovaSeq6000 on ribo- and globin depleted RNA samples originally collected from PaxGene Tubes. Analysis was conducted jointly by TGen and the Institute of Translational Genomics at USC. The sequencing reads were mapped to genome assembly GRCh37 (hs37d5) using STAR [20] on genes specified in GENCODEv19. The number of mapped reads was quantified with FeatureCounts [21].

PPMI DaTscan SPECT imaging data measure the amount of dopamine transporter (uptake) in four regions of brain striatum (right/left caudate and putamen). DaTscan images were acquired at PPMI imaging centers following the PPMI imaging protocol. The collected images were sent to PPMI imaging core lab at Institute for Neurodegenerative Disorders (New Heaven, CT, USA), where potential deficits in dopamine uptake were assessed by experts and visual inspection and interpretation were provided. Raw SPECT projections were iteratively reconstructed on a HERMES workstation (HERMES Medical Solutions, Stockholm, Sweden) using the HOSEM program (HERMES OSEM, based on the original OSEM algorithm [22]), by each imaging center. The reconstructed data with HOSEM were transferred to PMOD (PMOD Technologies, Zurich, Switzerland) for additional preprocessing. The images underwent attenuation correction using the Chang 0 method [23]. Corrected data were then normalized to standard Montreal Neurologic Institute (MNI) space [24], in order that all scans shared the same anatomical alignment. The regions of interest (ROIs) were on right/left caudate and putamen and on the occipital cortex, which was used as the reference tissue. Striatal Binding Ratios for the 4 striatal ROIs were computed by extracting count densities in each region.

The PPMI structural imaging protocol includes a sagittal 3D T1-weighted MRI. Acquisition was at either 1.5T or 3T using either MP-RAGE (magnetization prepared rapid acquisition gradient echo) or SPGR (spoiled gradient) sequences: 170–200 slices; 1×1×1.2mm voxel resolution; 0mm slice gap; field of view = 256×256mm. All other parameters followed site-dependent manufacturer recommendations for the scanner used (GE/Siemens/Philips). Scans were read by a radiologist at each site to meet standards of clinical practice and ensure that there were no significant abnormalities. Cortical reconstruction and volumetric segmentation of MRI scans was performed using FreeSurfer-v6.0 software [25, 26]. This involves skull stripping, volumetric labelling, intensity normalization, grey/white matter segmentation and registration to established surface atlases. Cortical thickness was calculated as the closest distance from the grey–white matter boundary to the grey matter–CSF boundary at each vertex.

In our study, we considered all data collected at the time of first visit (baseline). This choice allows temporal coherence among the different types of data used and avoids potential biases due to data collection time points and potential disease progression.

### 2.2 Genotyping data preparation

GWA studies search for statistical associations between genotype and phenotype, by testing thousands and even millions of SNPs for relations with the studied traits [27, 28]. Often, raw genotyping data contain errors, which could consistently affect the final outcome of GWAS. These errors can be originated from different sources, such as contaminations, poor quality of DNA samples or poor hybridization on the genotyping array [29]. Therefore, it is necessary to perform a quality control (QC) step, to clean genotyping data.

Even though some statistical analysis underlying QC and GWA studies could be carried out with software packages in R or Python, it is more convenient to use frameworks specifically tailored for genetic data analysis. Currently there are several comprehensive tools specifically designed for GWAS analyses, such as PLINK [30], GenABEL [31], SNPTest [32], or MaCH [33]. PLINK toolkit provides a comprehensive framework for genetic data management, quality control and association tests. Since PPMI genotyping data were provided in PLINK format, we used PLINK toolkit (version 1.9b) to perform QC and test for SNP-trait associations. However, similar analyses could be carried out using other GWAS toolkits.

In our study, we prepared genotyping data by performing extensive QC steps. First, we merged ImmunoChip and NeuroX in a single dataset. We then excluded from the study those SNPs showing genotype missingness rate > 5%, minor allele frequency (MAF) < 1% and with Hardy-Weinberg equilibrium < 1 · 10 ^−6^. Since most PD-related genetic variants are located on autosomal chromosomes [34], we excluded SNPs occurring in sexual and mitochondrial chromosomes. After QC remained 164,548 SNPs for further analysis. Moreover, we adjusted data for population stratification using Principal Component Analysis (PCA). Then, we kept in our study 260 (HC = 87, PD = 173) unrelated subjects with European ancestry, for which were available complete genotyping, neuroimaging and transcriptomic data.

### 2.3 Neuroimaging data preparation

Often, raw neuroimaging data values contain outliers, which can significantly drive SNP-traits statistical associations, introducing potential bias in the results. Therefore, to reduce such effects it is common practice to normalize imaging data. In our case study, we normalized both DaTscan and MRI data applying rank-based inverse normal transformation (r-INT) [35]. The main advantage of r-INT is that it can manage outliers without affecting the overall normalization procedure.

### 2.4 Preparation of transcriptomic data

Transcriptomic data are commonly used to compare and associate gene expression profiles with phenotypic traits, among different samples through differential expression analyses (DE). Currently, there are several comprehensive computational frameworks to manage and analyze RNA-seq data through the application of machine learning and statistical methods, such as Limma+Voom, EdgeR, DESeq2, NOIseq or SAMseq [36–41]. In our analysis we used DESeq2, since it has been shown to be one of the best frameworks in terms of precision, sensitivity and accuracy [42].

Often, transcriptomic data contain potential problems, such as low read counts or sample outliers, which can introduce errors during the analysis. Therefore, QC is fundamental to remove such issues from RNA-seq data.

It is common practice to filter RNA-seq data for highly expressed genes, for example, removing genes not expressed in all samples, or removing those not expressed in at least one sample for each condition group analyzed. In our analysis we applied the latter. Another important QC step is to check for potential outliers or batch effects determined by external features. For our analysis we considered 260 unrelated subjects with European ancestry, with complete genotyping, neuroimaging, and transcriptomic data.

### 2.5 Multi-view methods for imaging genetics analysis

Most of GWA studies on complex diseases focused on statistical association between SNP and single phenotypes. However, several studies showed that one genetic variant can affect different traits (pleiotropy), particularly in complex diseases [43, 44]. Therefore, considering single phenotypic traits could result in a loss of statistical power in the identification of genetic mechanisms underlying complex diseases. Instead, taking into account multiple correlated phenotypes can improve the discovery of genetic variants, which could influence different traits underlying the onset of the studied complex disease [45], and can provide new biological insights revealing pleiotropic effects [46].

In recent years, several methods have been proposed to analyze multivariate phenotypes [47]. These methods employ two main approaches (views): computing summary statistics from univariate analysis, or providing generalized models combining phenotypic measures used to test for variant-trait associations.

Significant variant-trait associations found during GWAS analysis can provide new biological insights on genetic mechanisms underlying the studied trait. Therefore, GWAS results are functionally annotated and often validated, for example by mining databases providing genes observed to be affected by variants’ presence. When available, transcriptomic data can be exploited to provide further and more accurate functional annotation and validation to GWAS results. In fact, transcriptomic data can be used to confirm, confute and even add hypotheses on functional consequences of variants associated with the studied traits.

Below, we describe our multi view-based workflow, composed of three main phases: *Individual view, Integrated view* and *Functional interpretation* (see **Figure 1**). It covers the key steps of an *imaging genetics* analysis, combining genetic, brain imaging and transcriptomic data.

**Figure 1.**
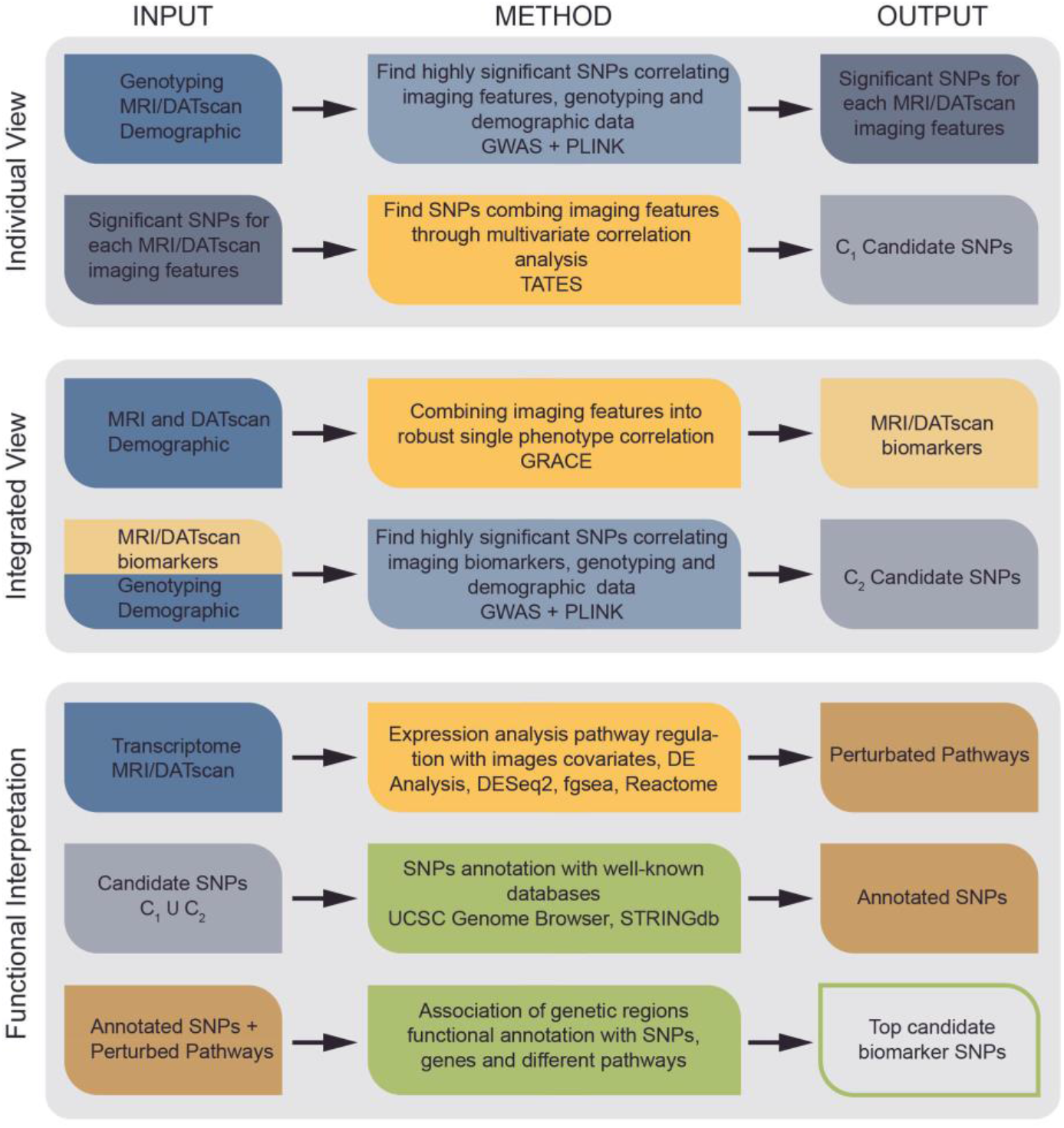
Workflow overview. The proposed workflow for imaging genetics is divided in three main branches: *Individual view, Integrated view* and *Functional interpretation*. The *Individual view* searches for SNP-trait associations by considering individually each neuroimaging feature and combines each single result after the GWAS analysis. The *Integrated view* phase aggregates neuroimaging data in a single measure, which is used as phenotype during the GWAS analysis. The *Functional interpretation* step aims to combine and annotate the potential genetic biomarkers found during the two previous phases. The candidate SNPs retained after *Functional interpretation* are returned as potential biomarkers describing the studied traits.

#### 2.5.1 Individual View

GWA studies considering multivariate phenotypes can improve the discovery of pleiotropic variant effects, providing new perspectives on complex diseases. A widely used strategy exploiting multiple traits potential is to combine individual test statistics in a single comprehensive measure [48]. In fact, several methods have been proposed to integrate single univariate test results in single summary statistics, such as TATES, Fisher Combination (FC), or Adaptive Fisher Combination (AFC) [49–51].

The main idea underlying these methods is to combine variant-trait association *p*-values obtained in standard univariate GWA studies, while correcting or weighting *p*-values according to phenotype correlation, for example. The Trait-based Association Test that uses Extended Simes procedure (TATES) [49] is a widely used method employing this approach to combine univariate statistics. TATES detects effects across correlated traits within the same group of subjects. It combines *p*-values of individual GWAS in single comprehensive statistics, by applying weights to account for correlations between the tested phenotypes and their number. Other interesting proposals are FC [50] and AFC [51] methods. Briefly, univariate GWAS *p*-values are combined through a permutation procedure, correcting for phenotype correlation. In our case study functional and structural brain images were collected from the same set of subjects. In such a setting TATES has been suggested as an efficient choice to model potential pleiotropic effects [52].

During the *Individual view* step (see **Figure 1**) of the proposed imaging genetics protocol is implemented the multivariate GWAS performed combining individual univariate result statistics in a single statistical measure. Therefore, this step carries out a GWAS on each feature separately and the corresponding individual results are combined in a single comprehensive statistical measure. In our case study on PD, we performed SNP-trait association tests with PLINK. PLINK computes linear regression models using variants as predictors and trait values as response. Moreover, PLINK regression analysis enables the use of covariates to correct the model response.

In our case study we modulated the response by correcting the models for individuals age, sex, education years and the first five principal components (PCs) of the relatedness matrix to account for population structure. Only when considering MRI phenotypes, we adjusted the models also for intracranial volume. Based on the linear regression model, PLINK provides a *p*-value assessing the statistical significance of SNP-trait association for each polymorphism. The individual univariate results were combined with TATES, which provided a single comprehensive statistic assessing SNP-phenotype association for each polymorphism.

#### 2.5.2 Integrated View

GWA studies can exploit multivariate traits to find potential pleiotropic variants, by computing generalized models combining phenotypes. These models are then used to test potential effects of variants on different phenotypes. Therefore, GWAS are performed searching statistical associations between SNPs and model measures describing the combined phenotypic traits. During the last decade, have been proposed several methods and tools to compute generalized models summarizing quantitative traits for GWAS, such as CCA, Multiphen or MANOVA [53–55]. Canonical correlation analysis (CCA) [53] and Multiphen [54] summarize different quantitative phenotypes by computing the statistical significance of traits most associated with each variant against the null-hypothesis of no association.

The main difference among the two methods rely in genotyping distribution: CCA assumes genotypes as normally distributed, while Multiphen as ordinal. In GWA studies MANOVA [55] tests how each SNP describes the variance present in multiple quantitative phenotypic traits, extending the univariate procedure (ANOVA) to multivariate analyses. The general idea behind this GWAS approach is to compute comprehensive generalized models or measures combining different phenotypes. Thus, also other methods computing generalized models focused on specific problems can be employed, such as modeling trait heritability [56] or brain imaging phenotypic changes over time [57–59].

During the *integrated view* phase (see **Figure 1**) of the proposed workflow we employed this approach for multivariate GWAS in order to reinforce the set of variants potentially affecting the considered traits. In our case study, we combined DaTscan and MRI traits by imaging modality using GRACE [57]. GRACE computes models to describe phenotype evolution at different time points, by estimating long-term growth curves fitting values changes over time. To summarize our phenotypic data, we constructed growth curves with GRACE as a function of the patients’ age. Therefore, GRACE assigns to each subject a quantitative score, measuring how much the associated traits are far from the expected growth curve.

During the GWAS analyses, we used PLINK to test for potential statistical associations between SNPs and model scores. We performed the GWAS analyses by correcting for age, sex, education years and the first five PCs of the relatedness matrix. For MRI traits we corrected the model also for intracranial volume. The GWAS provided for each variant a *p*-value assessing how the SNP affects the combined phenotypes.

#### 2.5.3 Functional interpretation

Differential expression analysis helps prioritizing differences in gene expression patterns linked with phenotypic traits, providing an insight on functional level. Therefore, the differences observed in gene expression profiles can be used to estimate through enrichment analysis the potential changes in biological pathways correlated with the study traits.

Enrichment analysis focuses on retrieving the biological pathways in which are functionally involved sets of genes. Several databases like GO, KEGG and Reactome [60–65] provide experimentally validated pathways and the involved genes. Enrichment methods can be classified in three main families [66]: over-representation analysis (ORA), functional class scoring (FCS) and pathway-topology (PT) methods. Briefly, ORA methods recover enriched pathways through overrepresentation analysis, which compares the proportion of involved annotated genes with the input list, used in Onto-Express [67,68] and GenMAPPP [69,70]. FCS enrichment returns functional pathways by considering gene-level statistics or LogFold changes of the differential expressed genes retrieved during DE analysis, used by GSEA [71,72] and SAFE [73]. PT enrichment methods, instead, extend FCS and ORA approaches integrating the previous algorithms with the analysis of pathways network topology, used by SPIA [74] and NetGSA [75].

During the *Functional Interpretation* step we performed DE and enrichment analysis to confirm and annotate the biological relevance of the results obtained during *Individual* and *Integrated views*. This step focuses on analyzing transcriptomic data to retrieve changes in expression patterns which could be potentially associated with imaging values fluctuations. We performed DE analysis with DESeq2 [38] computing a statistical model adjusted by age, sex and education year. Based on the statistical model, DESeq2 provides for each gene the LogFold change and its statistical significance (*p*-value and adjusted *p*-value). The statistically significant genes were then used to infer potential biological meanings to the results through enrichment analysis. We performed enrichment employing the fgsea algorithm [76], a fast and accurate implementation of the GSEA[71,72] and Reactome[65] database which contains several peer-reviewed and manually annotated biological pathways.

There are several approaches to infer functional annotation for SNPs retrieved during *Individual* and *Integrated view* steps. GWAS results can be directly annotated by specifically tailored software, such as FUMA [77], which infer the potential genes and biological pathways affected by significant genetic variants. Genes potentially regulated by GWAS SNPs can be also manually retrieved from databases such as GTEx [78,79], a curated public resource that studies potential correlations between gene expression patterns and regulatory SNPs. Otherwise, genomic sequences can be inspected with dedicated tools such as UCSC Genome Browser [80] searching for genes neighbouring the SNPs recovered during *Individual* and *Integrated view* phases. Moreover, by recovering Protein-Protein Interaction networks (PPI) of genes influenced by significant genetic variants, can be also captured potential distal consequences providing a comprehensive perspective of the biological processes affected. PPI can be retrieved from many curated databases such as STRING [81] or MINT [82]. In our case study on PD, we recovered the influenced genes and their interactive partners by using UCSC Genome Browser and STRING database.

## 3 Results

To showcase the key steps of an *imaging genetics* analysis and assess the proposed multi-view analysis protocol, we tested the method to search for potential genetic biomarkers using genotyping, neuroimaging and transcriptomic data on PD from PPMI, as described in Sections 2.1–2.4.

We investigated DaTscan Caudate and Putamen imaging to search for potential genetic biomarkers linked to dopamine uptake loss observed in PD subjects. Similarly, we explored parahippocampal structural MRI images [83] for potential genetic factors associated with cerebral atrophy and cognitive impairment, typically observed in PD patients.

Performing the *Individual view* step (see Section 2.6.1), we observed that GWAS analyses returned 5 statistically significant genetic variants (DaTscan = 3, MRI = 2) associated with the studied phenotypic traits (see **Figure 2**). These SNPs could constitute potential genetic biomarkers describing the differences in DaTscan and MRI values between healthy and PD subjects. We defined as potential biomarkers those variants showing statistically significant SNP-trait association (Empirical *P*-value < 1 · 10^−5^). Interestingly, we observed that the variant rs79761481, recovered employing DaTscan data, was located within the CD6 gene. This gene belongs to the CD genes family, which have been associated with PD onset at immune system level [84]. The remaining variants recovered during the *Individual view* phase were mapped in intergenic regions.

**Figure 2.**
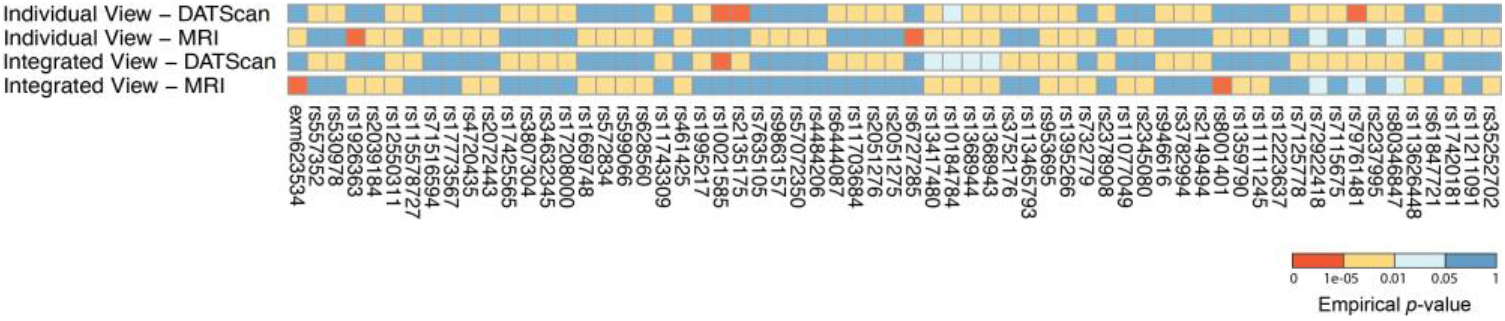
Individual and Integrated View resulting SNPs. Resulting significant SNPs after applying the Individual and Integrated view phases. The SNPs are colored by empirical p-value values.

Similarly, in *Integrated View* step (see Section 2.6.2) GWASs returned 3 additional potential genetic biomarkers (DaTscan = 1, MRI = 2) (see **Figure 2**). Again, we consider as potential biomarkers those variants showing statistically significant SNP-trait association (Empirical *P*-value < 1 · 10^−5^). We found that *exm623534* variant mapped within the sequence of ERV3-1 gene, while the remaining SNPs were mapped within intergenic regions.

We highlight the different significant SNPs obtained by using the proposed workflow that integrates different analysis flows (individual and integrated views) and heterogeneous data, to recover potential genetic biomarkers.

To functionally annotate and validate the findings of *Individual* and *Integrated View* steps, we performed the *Functional Interpretation* phase of the workflow as described in Section 2.6.3. During this step we performed DE and Enrichment analyses to inspect the differences in gene expression profiles and explore the biological perturbed pathways potentially linked to DaTscan and MRI values fluctuations. Not significant differentially expressed genes were filtered out keeping only those with adjusted *p*-value lower than 0.1. The results obtained highlight several differentially expressed genes for each considered imaging features analyzed (DaTscan and MRI). Moreover, from the functional enrichment analysis, after filtering for statistically significant pathways (adjusted *p*-value < 0.1), we retrieved several potentially perturbed gene sets associated with neuronal system, signal transduction, immune system, cell cycle and metabolism (see **Figure 3**) reinforcing the results quality and highlighting the biomarkers relevance, obtained by combining different imaging data type to genetics. The candidate biomarkers resulting from associating DaTscan are involved in a larger number of significant pathways compared to the ones obtained using MRI. Surprisingly, performing enrichment on the PPI genes retrieved and comparing the results with those obtained from the transcriptomic data analysis, we were able to retrieve common pathways potentially involved in the disease onset and progression such as Innate Immune System, Neutrophil degranulation and TCR signaling signaling pathways [85–87].

**Figure 3.**
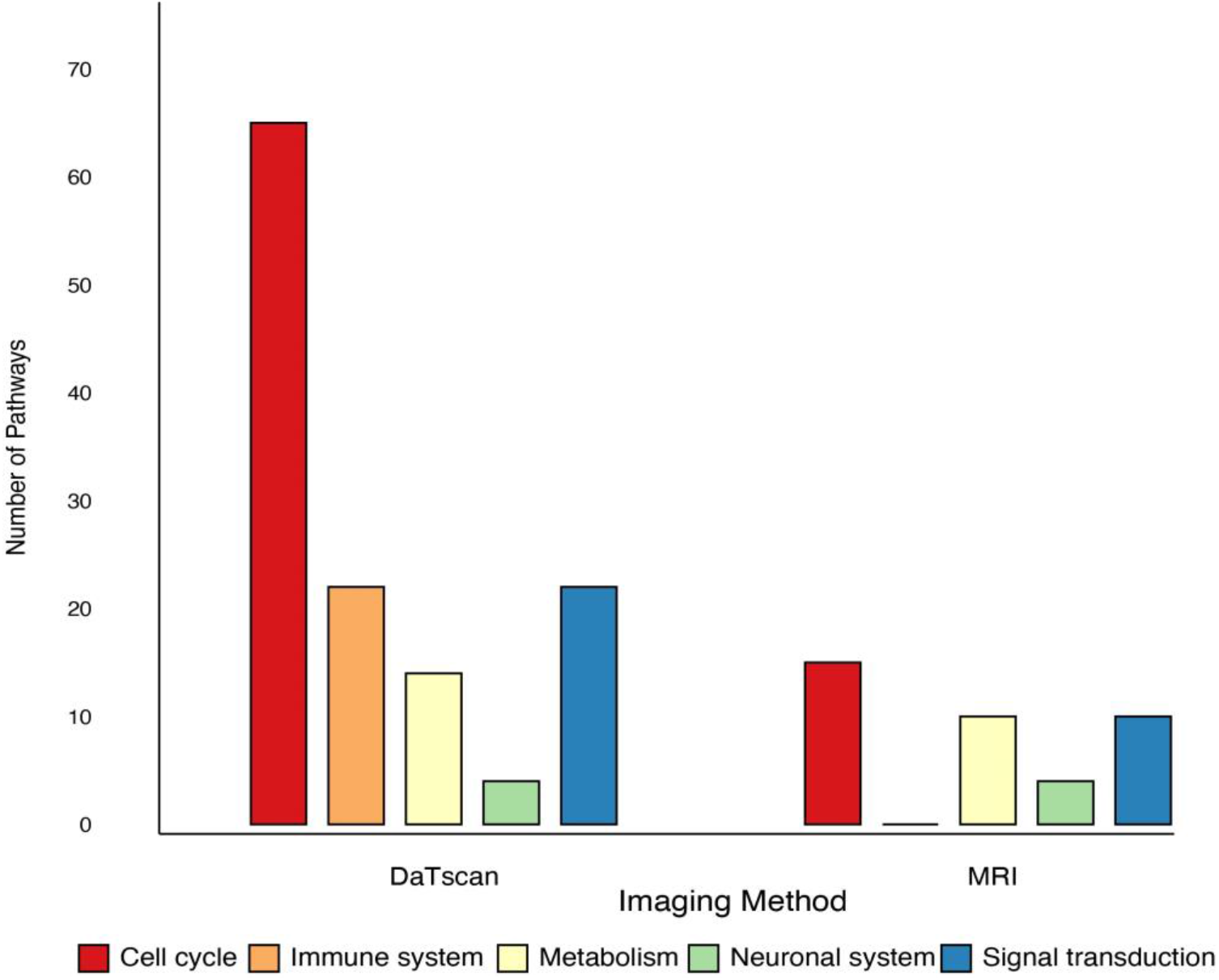
GSEA pathways retrieved from transcriptomic data analysis: barplot showing the number of REACTOME pathways, retrieved with fgsea package, for the imaging features on which the enrichment was performed. Pathways are grouped by macro biological function into five different sets including: *Cell Cycle, Immune system, Metabolism, Neuronal system, Signal transduction*.

## 4 Conclusions

We review the steps of an *imaging genetics* analysis by investigating PD-related data, by presenting a multi-view based workflow to analyze genetic and neuroimaging data. Unlike previous works, we studied dopamine uptake loss and cognitive impairment in PD, by searching for association among SNPs, dopamine uptake levels and brain structural variation. The proposed method can be a valid option with respect to a classical univariate analysis. With our workflow we are able to retrieve interesting genetic variants which could potentially play a role in PD onset and development.

## Acknowledgements

Data used in the preparation of this article were obtained from the Parkinson’s Progression Markers Initiative (PPMI) database (www.ppmi-info.org/data). For up-to-date information on the study, visit www.ppmi-info.org. PPMI – a public-private partnership – is funded by the Michael J. Fox Foundation for Parkinson’s Research funding partners 4D Pharma, Abbvie, Acurex Therapeutics, Allergan, Amathus Therapeutics, ASAP, Avid Radiopharmaceuticals, Bial Biotech, Biogen, BioLegend, Bristol-Myers Squibb, Calico, Celgene, Dacapo Brain Science, Denali, The Edmond J. Safra Foundaiton, GE Healthcare, Genentech, GlaxoSmithKline, Golub Capital, Handl Therapeutics, Insitro, Janssen Neuroscience, Lilly, Lundbeck, Merck, Meso Scale Discovery, Neurocrine Biosciences, Pfizer, Piramal, Prevail, Roche, Sanofi Genzyme, Servier, Takeda, Teva, UCB, Verily, and Voyager Therapeutics.

## Key points

- We review the steps of an *imaging genetics* analysis by investigating Parkinson PD-related data, collected from PPMI data portal.
- We review the analysis steps in action by implementing an imaging-genetic workflow, integrating heterogeneous data (genetic, imaging and transcriptomics data).
- The analysis investigates and combines different methods to analyze the data.
- We show that our analysis using the multi-view workflow was able to reach reliable results for PD data.
- The source code of the implemented multi view based method is freely distributed to the communities.

## Data availability

The code underlying this article is available at https://github.com/InfOmics/MUVIG. Data is accessible through the PPMI data portal, at https://www.ppmi-info.org.

